# Lineage: Visualizing Multivariate Clinical Data in Genealogy Graphs

**DOI:** 10.1101/128579

**Authors:** Carolina Nobre, Nils Gehlenborg, Hilary Coon, Alexander Lex

## Abstract

The majority of diseases that are a significant challenge for public and individual heath are caused by a combination of hereditary and environmental factors. In this paper we introduce Lineage, a novel visual analysis tool designed to support domain experts who study such multifactorial diseases in the context of genealogies. Incorporating familial relationships between cases with other data can provide insights into shared genomic variants and shared environmental exposures that may be implicated in such diseases. We introduce a data and task abstraction, and argue that the problem of analyzing such diseases based on genealogical, clinical, and genetic data can be mapped to a multivariate graph visualization problem. The main contribution of our design study is a novel visual representation for tree-like, multivariate graphs, which we apply to genealogies and clinical data about the individuals in these families. We introduce data-driven aggregation methods to scale to multiple families. By designing the genealogy graph layout to align with a tabular view, we are able to incorporate extensive, multivariate attributes in the analysis of the genealogy without cluttering the graph. We validate our designs by conducting case studies with our domain collaborators.

## 1 INTRODUCTION

**S**TUDYING ancestry and familial relationships, i.e., genealogies, is both a pasttime enjoyed by amateurs and a research area for professionals [53]. It is hence not surprising that there are numerous tools to record and visualize genealogies. Yet, most of these tools focus on analyzing family structures for historical purposes, and only a few target a clinical use case of analyzing genealogies in the context of complex, hereditary diseases. Geneticists, on the other hand, have long used genealogical graphs to study how a genetic disease manifests itself in families. They use drawing conventions and standardized symbols to show both the family structure and the phenotype, i.e., the observable characteristics of an individual [6]. These charts can provide insights about the heritability and segregation patterns of genetic diseases. In their current form, however, they are predominantly useful for Mendelian diseases, or genetic diseases caused by a small number of mutations. Complex diseases such as cancer, autism, diabetes, obesity, and psychiatric conditions such as depression or suicide, are known to have hereditary components that are regulated by a multitude of genes, each having a modest effect on risk, and also to depend strongly on environmental conditions and chance. When studying these conditions in a population, it is imperative to simultaneously consider genetic similarities, shared characteristics of the phenotype, and environmental conditions. Also, for these polygenic conditions, one needs to consider significantly larger populations to reason about hereditary relationships and pursue the discovery of genetic risk mutations.

Current medical or historical genealogy visualization tools are ill equipped to help researchers find patterns in these large, highly multivariate graphs of families and their rich medical histories. In this paper, we present a novel genealogy visualization tool that we have developed in collaboration with psychiatrists and geneticists studying the genetic underpinnings and the environmental factors of suicide and autism. We use data from the Utah Population Database^1^, a uniquely rich resource for population-based analysis of hereditary diseases.

We contribute a novel technique to visualize large, treelike graphs (rooted, directed graphs that have some cycles but are predominantly in tree form) associated with rich numerical, categorical, and textual attributes. Our approach leverages the treelike structure of the graphs to produce a linearized layout that enables the direct association of the nodes with rich attributes in a tightly integrated tabular visualization. We address the issue of scalability by introducing novel forms of degree-of-interest-based aggregation that preserve the structure of the graph, and, if desired, also provide an overview of the attributes of aggregated individuals. We demonstrate our technique in the context of genealogical data, and we argue that it can be equally applied to other multivariate trees or tree-like graphs.

We also contribute a detailed characterization of the domain problems and the domain data as they are encountered when analyzing large, clinical genealogies^2^ and a set of task and data abstractions derived from these characterizations. Finally, we contribute the open-source Lineage visualization tool (https://lineage.caleydoapp.org), shown in Figure 1, which implements the technique, and describe multiple design decisions tailored to genealogical data visualization.

**Fig. 1.**
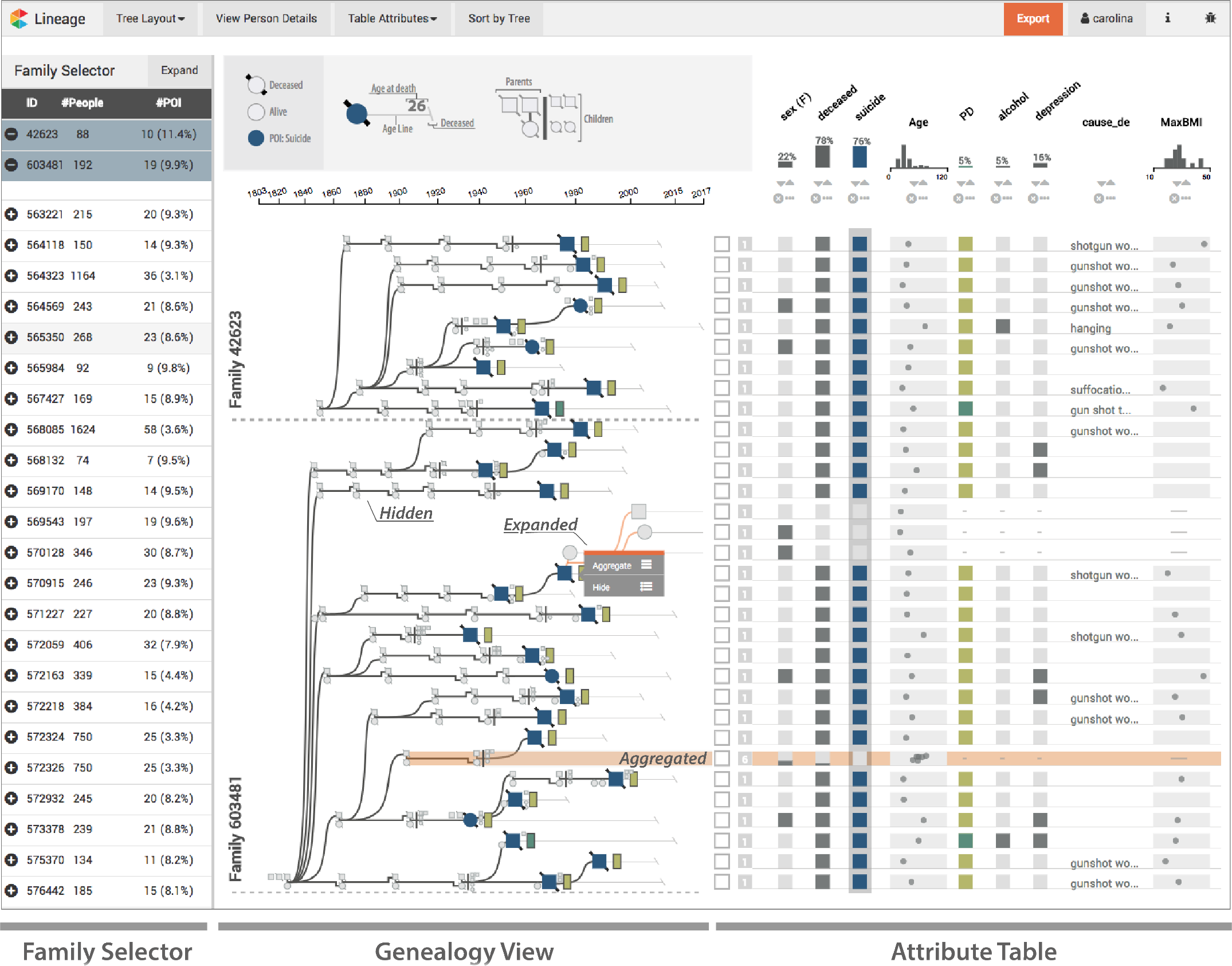
Lineage visualizing the genealogy of two families with increased numbers of suicides. The genealogy view shows the family relationships in a linear tree layout, where each node corresponds to a row in the associated table. Suicide cases are highlighted in blue, and a glyph next to the nodes indicates whether individuals were diagnosed with a personality disorder. The branches use different levels of aggregation (hidden, aggregated, expanded). The table shows detailed attributes about individuals, or, when branches are aggregated, for groups of individuals.

Lineage is in the process of being adopted by our collaborators, and has undergone iterative design refinements. We have also demonstrated it to other research groups working with genealogical and genetic data and have encountered overwhelming enthusiasm. We validate this work in an illustrative usage scenario and through qualitative user feedback from domain experts.

## 2 DOMAIN BACKGROUND AND DATA

Our collaborators study the genetic underpinnings and the environmental factors influencing psychiatric conditions, such as autism and suicide, using detailed genealogical, clinical, and genetic data. In this paper, we will focus on suicide, yet our methods are easily transferable to other complex, multifactorial conditions and diseases. Suicide is a high-impact application, as it is one of the leading causes of life-years lost [76] and the 10th most common cause of death in the United States [56]. Suicide is believed to be caused by a complex combination of risk factors, including environmental stressors and genetic vulnerability. Aggregated data across multiple large studies has produced a heritability estimate of completed suicide of 45% [51], [61]. Genetic risk factors for suicide are complex and can be classified as multiple subtypes. These subtypes often are characterized by co-occurring psychiatric conditions (comorbidities) and/or a combined risk of psychiatric diagnosis. For example, the genetic risk for schizophrenia is also associated with a risk for suicide [68].

Our collaborators have compiled a unique dataset of suicide cases, including DNA and clinical profiles on 4,017 cases. These cases are linked to the Utah Population Database (UPDB), which provides genealogical data. Genealogies describe the familial relationships of individuals across multiple generations.

Figure 2 shows two genealogies using standardized drawing conventions [6]. Females are drawn as circles, males as squares. Couples are connected by an edge, and children connect to this edge using orthogonally routed links. The vertical position of nodes is given by their generation. A phenotype of interest is marked by a filled-in node. When studying family relationships, a common approach is to draw family trees considering the ancestry of an individual. Figure 2(a), for example, shows the family of the woman marked in black. The genealogy includes her two siblings and traces her family tree for two generations to include parents, uncles and aunts, and grandparents.

**Fig. 2.**
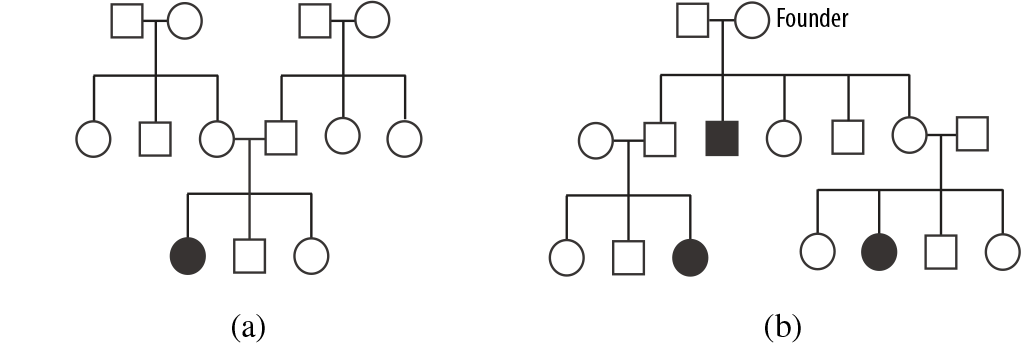
Two genealogies using standardized symbols focusing on different aspects of the family structure. Females are shown as circles, males as squares. Individuals with a phenotype of interest are filled-in in black. (a) A genealogy showing the family of the female in black, including siblings, parents, uncles/aunts, and grandparents. (b) A genealogy based on a founder, tracing down generations to include the families of individuals with a phenotype of interest (black).

In contrast, our collaborators are interested in understanding genetic relationships between individuals afflicted with a condition and hence care about individuals who share genetic variants. They select families for study that have a statistically increased rate of a condition. These family trees are constructed by tracing cases back to a “founder”, as illustrated in Figure 2(b). The underlying hypothesis is that such founders have genetic risk variants that they passed on to their descendants. Within the genealogy, the likelihood of genetic homogeneity is increased, and is more easily detected through the repeated occurrence of the genetic risk variant in the familial cases. Note that this genealogy contains only individuals who are descendants of one founder and his or her spouse, with the exception of spouses of descendants. Also, the dataset contains only individuals with direct links to a case; i.e., siblings, descendants, and direct ancestors are included, whereas, for example, uncles/aunts and cousins are not.

The dataset our collaborators have compiled contains about 19,000 suicide cases, including 4,585 recent cases with detailed data, backed by family structures made up of 118,000 individuals from 551 families. Suicide is frequently associated with psychiatric comorbidities, i.e., co-occurring chronic conditions, such as depression, bipolar disorder, substance abuse, PTSD, or schizophrenia [68]. Also, nonpsychiatric conditions such as asthma [30] may play a role in some cases. Environmental factors, such as socioeconomic status, pollution, and seasonality, are also known to be factors in suicide [2]. To capture this information, our datasets include demographic variables such as gender, race, age at death, method of death, family demographics (marriage, divorce, number of siblings/children), and place of residence at the time of death. The datasets also include records of other diagnoses captured as codes from the International Classification of Diseases (ICD) systems, the frequency with which these diagnoses were made, and the time of the first diagnosis.

To summarize, each of our many graphs describes a family, with individuals as nodes and family relationships as edges. Since the graphs are constructed by tracing ancestry to a founder, they are predominantly tree-like, but they do include cycles, for example, when two cousins have offspring. In addition, we have attributes on the individuals/nodes in the graphs of various data types, including numerical, categorical, temporal, geographic, and textual data. These attributes are often sparse because only about 10% of individuals in the dataset have committed suicide, and our detailed records extend to only about 2% (4,017) of individuals across all families. These detailed records capture about 3,000 dimensions that contain demographic information and information about the manner of death, but predominantly contain comorbidities in the form of disease codes and the time and frequency of these diagnosis. These dimensions are themselves often sparse because, among other reasons, a colloquial diagnosis such as “depression” can be recorded using one of about 30 ICD codes.

## 3 DOMAIN GOALS AND TASKS

This project is rooted in a collaboration with faculty, clinicians, analysts, and graduate students in the Department of Psychiatry at the University of Utah. Six domain experts participated in the project. We loosely followed the design study methodology proposed by Sedlmair et al. [64]. Our “discover” phase consisted of multiple meetings with individual collaborators and with the whole group as a team, studying the domain literature and the tools they currently use. We also ran a creativity workshop, specifically the *wishful thinking* component described by Goodwin et al. [24], involving all the collaborators. In the workshop, we asked participants to think about the analysis of suicide data and then discuss in small groups and take notes on post-its about what it is they would like to *know, see*, and *do*. This idea-generation phase was followed by a phase in which the teams had to prioritize their insights and then finally give the whole team an overview of their key ideas. We recorded the workshop and transcribed both the audio and the post-its. We then coded the artifacts and three themes emerged: the *data*, the *factors involved in suicide*, and the *analysis tasks*. The insights on the data and the factors involved in suicide were described in the previous section.

The overarching goal of our collaborators is to gain a better understanding of the determining or associated factors of suicide. Our collaborators classify the factors associated with suicide into comorbidities and demographic, genetic, and environmental factors. Specifically, they are interested in identifying and defining detailed phenotypes associated with suicide and the degree to which these phenotypes are familial. By finding people who are similar to each other in a relevant way, our collaborators hope to reason about genetic homogeneity, i.e., shared genetic factors contributing to suicide. They currently rely only on familial structure as a proxy for genetic homogeneity. However, they recognize that this approach is limited both as too broad — it is possible that they should consider only a part of a family — and as too narrow — people outside a family who have a similar phenotype could also have a similar genotype. Robust and detailed phenotypes are, of course, also interesting by themselves, because they can be used, for example, as part of a risk assessment in a clinical context.

It is important to note that the contextual knowledge of a researcher is beneficial to the task of classifying a phenotype. For example, a diagnosis of depression is weighted differently if it is diagnosed dozens of times and was first diagnosed early in a patient’s life. Similarly, a young person who commits suicide in a rural community is unlikely to have a detailed medical history. Hence, such a case could be similar to others, even if certain phenotypes are not recorded, if other factors, such as a close familial relationship, indicate it.

Our collaborators need a visualization tool that is embedded in a larger analysis process, one that includes calculating statistically significant familial risk (upstream) and searching for shared genetic variants (downstream). They need a tool that focuses on finding individuals and families that are “interesting” with respect to both their relatedness and their attributes, which can then be used in further analysis and validation.

We identified the following domain tasks as the most important aspects in the workflow of our collaborators:

### T1 Select families of interest

The analysts want to select a family by browsing, using prior knowledge, or employing a data-driven approach. The analysts want to select a family either by browsing, or by selecting a specific family based on prior knowledge, or in a data driven way. An example of the data-driven approach is to find families with high rates of suicide or with individuals for which suicide co-occurs with bipolar disorder.

### T2 Analyze individual case

Our collaborators need to investigate the context of a case. For example, a potential genetic component contributing to suicide is judged differently if the person had many psychiatric comorbidities and committed suicide at a young age, compared to a late-life suicide of a person with a terminal disease.

### T3 Compare cases

This task encompasses comparing individuals and identifying shared attributes to characterize a potentially meaningful shared phenotype. It also pertains to analyzing how the individuals are related, which can indicate the likelihood of shared genetic traits. Insights on shared environmental factors can be gleaned from both the family structure and the attributes. For example, siblings are likely to be exposed to the same environment in their childhood, whereas cousins might not. Similarly, two people living in the same area are potentially of similar socioeconomic status.

### T4 Judge prevalence and clusters of phenotype

The number of suicide cases and the prevalence of comorbidities vary greatly between families and between branches of a single family. Judging how common a phenotype is in a family or in part of the family is helpful in identifying subsets of interest for further study.

### T5 Compare families

Once an interesting observation has been made in one family, our collaborators want to be able to investigate whether similar cases also appear in other families. For example, when an association of asthma with suicide is discovered, they want to know whether it is isolated in one family or occurs in multiple families and/or individuals.

### T6 Quality control

Although not an analysis task per se, our collaborators also need to judge the quality of the data and report errors back to the central database. A common data error we have seen is disconnected components or detached nodes, which are caused by missing information about an individual’s mother or father.

Most of these domain tasks rely on both studying the topology of the network, i.e., the family relationships, and investigating the attributes associated with the individuals. For example, the “compare cases” task (**T3**) relies on both the graph and the attributes to, for example, reject an outlier in an otherwise well-defined phenotype within a family, if that outlier is only distantly related to other cases.

## 4 RELATED WORK

We focus our discussion of previous work on specialized genealogy visualization tools and on multivariate network visualization, since genealogies are highly multivariate graphs. With regard to multivariate network visualization approaches, we also restrict our discussion to explicit layouts (i.e., node link layouts) because implicit layouts (such as SunBursts and treemaps) are ill suited to visualize attributes at all levels of the hierarchy; and matrices are not an ideal choice for genealogies since (1) the nodes are only sparsely connected, hence wasting space, and (2) matrices are ill suited for path tracing, which is a common task of our collaborators.

### 4.1 Multivariate Networks

A multivariate network is a graph where the nodes and the edges are associated with potentially rich attributes [32]. Many
graph visualization techniques are optimized for either topology or attribute-based tasks [72], yet in many applications topology and attributes have to be judged in concert [59]. When analyzing genealogies, for example, our collaborators want to understand how two people with a similar phenotype are related, requiring them to first identify the phenotypes using the attributes, and then to judge their relatedness using the topology of the genealogy.

Partl et al. [59] classify four basic approaches to visualize multivariate networks for explicit graph layouts: (1) on-node mapping, i.e., visualizing the attributes by changing a visual channel of the node mark or by embedding a small visualization in the node; (2) small multiples, i.e., showing the same graph multiple times and visualizing a different attribute on top of each small network; (3) separate, linked views for the graph and the attributes; and (4) adapting the graph layout to better fit the needs of attribute visualization.

These approaches have different strengths and weaknesses with respect to the tasks they enable. Lee et al. [41] distinguish, among others, topology-based tasks, i.e., tasks that are related to the network’s connectivity, and attribute-based tasks, i.e., tasks that are related to the attributes associated with the nodes.

Although **on-node mapping** simultaneously supports topology and attribute-based tasks, it does so for only a few attributes because the node size limits how many attributes can be encoded. Also, on-node visualizations are typically not aligned and have distractors between them, which makes accurate comparison difficult [13]. Gehlenborg et al. [22] review multiple systems that use on-node mapping for biological networks. An example for slightly more complex visualizations embedded on nodes is the Network Lens [29]. The work by van den Elzen and van Wijk [72] is a special case of an on-node mapping approach: instead of mapping data directly onto nodes in the networks, they aggregate nodes into supernodes, show the relationships between the supernodes, and visualize the attributes of these nodes in small, embedded visualizations.

Small multiples are also commonly used to visualize attributes on top of graphs. Barsky et al. [5] and Lex et al. [44], for example, use small multiples to show gene expression data on top of biological networks. Using small multiples for multivariate networks, however, has the disadvantage that the individual networks have to be rendered in less space, limiting their readability or the size of the graph for which they are useful.

Separate, linked views excel at visualizing the attributes and the graph individually, but do not support the integration of both well. Systems that use this approach [42], [66] rely on linking and brushing to associate a node with the representation of its attribute, which requires interaction to reveal relationships between the topology and attributes.

The fourth approach to multivariate graph visualization is to adapt the layout of the network so that the nodes can be easily associated with an effective attribute visualization. This approach is taken to the extreme in GraphDice [9], where nodes are positioned in a series of scatterplots based only on attribute values. Less extreme approaches are various linearization strategies where graphs are laid out such that associated attributes can be visualized in efficient tabular layouts, overcoming the drawbacks of completely separated linked views. Typically, trade-offs for optimizing the readability of the topology or the linear layout have to be made. Meyer et al. [52] manually linearize a complete network and render attributes next to the linear layout. This approach is efficient, but the complexity of the networks for which it is feasible is limited, and topological structures can be hard to see. Partl et al. [59] use interaction to extract paths from a network, linearize these paths, and associate the nodes in the paths with rows in a tabular visualization. This approach, however, requires interaction and works only for selected subsets of the graph. The recently published Pathfinder system [58] uses path queries on networks and presents the resulting paths in a linear, ranked list, juxtaposed with rich attribute data. This approach, however, is sensible only for tasks related to paths.

Our work falls into the category of adapting the layout by linearization. We leverage the fact that the genealogical graphs our collaborators are interested in are tree-like and linearize the positioning of the nodes in the tree. We use this tree to juxtapose scalable and perceptually efficient visualizations of the attributes.

### 4.2 Tree Visualization

Many examples of multivariate tree visualization techniques are available, yet none scale to more than a handful of attributes and work for both intermediate nodes and leaves at the same time. The on-node mapping strategies discussed in the previous section can also be applied to visualizations of trees (e.g., [12]), with the same limitations with respect to scalability. A common example for tree visualizations associated with many attributes is the use of a dendrogram tree derived from a hierarchical clustering algorithm that is aligned to a heat map [16], [43], [65]. Similarly, the leaves of evolutionary trees can be aligned to heat maps of the species’ traits [38], [39]. Engel et al. [17] use clustering to decompose a multidimensional dataset and represent it as a Structural Decomposition Tree. This approach is unique since it directly embeds a tree into a projection of a high-dimensional dataset, foregoing a tabular layout for the attributes. Also related to our approach are tree tables, since they can be found in file browsers, where the tree represents the structure of folder and files, and attributes such as the file size are shown. Tree tables generally do not provide aggregation functionality — a branch can be either collapsed or expanded, but cannot be aggregated.

Implicit tree visualization techniques such as tree maps [28], sun burst [69], or icicle plots [36] are well suited to visualize one or two attributes of nodes in trees (using size and color), but they do not scale to more attributes.

These approaches either scale to many attributes for the leaves of large trees, or are limited to a handful of attributes for all nodes of the tree. We are not aware of prior tree visualization approaches that also show rich attributes for intermediate nodes, either in aggregated form or for each node individually.

Our approach is also related to tree visualization techniques that provide dynamic aggregation, since we aggregate branches of trees to highlight nodes of interest. Our approach is based on the concept of degrees-of-interest functions [20], which is widely applied in trees [12], [27], including in the original paper, but is also related to other focus+context tree visualization approaches [55]. For a broad overview of other tree visualization techniques, we refer to the tree visualization reference by Schulz [63].

### 4.3 Genealogy Visualization

Genealogical charts, as shown in Figure 2, are widely used in genetic counseling and the literature on genetic diseases. They are well suited to visualize a single phenotype of interest, but they are not suitable to map a complex phenotype to the node. Our collaborators currently use Progeny [62], a commercial genealogy drawing tool that closely follows the standard for visualizing genealogies [6], [7]. (See the supplementary material for an example figure created with Progeny.) Although Progeny is well suited to draw these standard genealogies for use in presentations, it is ill suited for exploratory tasks, mainly because of its inability to efficiently encode attributes in the graph.

Interactive genealogy visualization tools that are designed to analyze disease clusters and to see disease propagation within families include PedVizApi [19], CraneFoot [46], Haploview [4], PediMap [73], and HaploPainter [70]. HaploPainter [70] visualizes genealogies and genetic recombination events below the individuals’ nodes. Although it shares the approach of showing metadata as rows associated with nodes with Lineage, it does not take a linearization approach to make values of different generations easy to compare, it does not aggregate the network, and it does not visualize different types of attributes. McGuffin and Balakrishnan [50] describe layout algorithms for complicated genealogical trees and introduce aggregation methods for subtrees, which we adopt.

Among tools that do not use the standard genealogical drawing conventions are Fan Charts [15], which uses the SunBurst technique to visualize genealogical trees, and the work by Mazeikla et al. [49], which employs a force-directed layout that considers similar phenotypes as additional attracting forces. Tuttle et al. [71] use an H-tree layout for scalable genealogy visualization, with the founder at the center and successive generations radiating out based on a fractal pattern. Ball [3] employs the idea to not represent generations as discrete units but use time to position the nodes, and also to draw a person’s life span. Kim et al. introduce TimeNets [34], a technique also focused on the temporal aspects of a genealogy. Although TimeNets is well suited to observe temporal changes in relationships between individuals, relationships between generations are harder to trace. The recent work by Fu et al. [18] focuses on visualizing the distribution of tree structures in many families. The tool combines a Sankey diagram showing properties of tree structures with explicit node-link diagrams on demand, but does not consider attributes of the nodes.

GenealogyVis [45] is a recent tool for visualizing genealogies to study historic data. Although it visualizes multivariate attributes, it addresses different needs — those of historians — and uses different approaches. Unlike in Lineage, attributes of individuals are not shown; rather, the focus is on demographic trends in (parts of) the network. Supplementary views, such as scatterplots and maps, allow historians to study, for example, migration patterns.

Genealogy visualization tools for animal genealogies face a different set of challenges compared to those for human genealogies, as the number of descendants sired by individual animals can be large, and complex interbreeding is common. Consequently, tree-based approaches are not well suited for these genealogies. Examples include CoVE [11] and VIPER [60]. VIPER introduces a sandwich view that CoVE also adopts. The sandwich view scales well to many descendants of an individual, but only explicitly encodes the relationships between parents and their children. More distant relationships can be revealed through highlighting. Helium [67] is a visualization technique for plant genealogies, which commonly have complex crossing. It uses color coding and scaling of nodes to encode up to two attributes.

GeneaQuilts [10] is a matrix-based technique where each row constitutes a person and each column a nuclear family. In early stages of our design process, we considered using a GeneaQuilt instead of our node link design, since GeneaQuilts produces a linearization of the graph that would be suitable for associating attributes. We ultimately decided against it because (1) the data we consider for the analysis of genetic relationship is predominantly tree-like, and hence, the complex design of GeneaQuilts that is necessary to accommodate general genealogical graphs is not justified for our simpler, tree-like datasets; (2) a key analysis task for the graph view is to judge the degree of relatedness between two nodes, which is not well supported by GeneaQuilts without interaction; and (3) our design for aggregation is more suitable for node-link diagrams.

A different approach to analyzing relatedness is to calculate “kinship coefficients” between individuals, i.e., to calculate path-based metrics for relatedness and visualize them in a matrix [33]. Although this approach is scalable, it is not suitable for reasoning about all patterns of inheritance.

A related tool that is concerned with visualizing phenotypes of patient cohorts is PhenoStacks by Glueck et al. [23]. PhenoStacks uses a tabular approach similar to what we use for our table.

## 5 VISUALIZING A MULTIVARIATE GRAPH

The tasks our collaborators need to address rely heavily on both the familial information contained in the genealogy graph, i.e., the topology, and the myriad of attributes associated with individuals (see Section 3). Of the strategies for linearization introduced in Section 4.1, only the linearization method enables an integrated analysis of topology and attribute at the scale of attributes we are interested in. However, none of the described linearization methods are suitable for the data and tasks of our collaborators. Here, we introduce a linearization method for tree-like graphs. We define tree-like graphs as rooted, directed graphs that contain cycles. The purpose of the linearization is to associate the nodes with rows in a table visualization.

A consequence of the linearization strategy is that the layout is not as compact as in other common layouts. To address this issue, we also introduce degree-of-interest-based aggregation strategies that integrate seamlessly with the linearized graph.

We illustrate this concept here using general, tree-like graphs, for now ignoring specific properties of genealogies. We later show in Section 6 that this approach extends to genealogies (where each person has two roots: their parents) with minor modifications, and also elaborate on design decisions we made that are specific to our data and application area.

### 5.1 Linearization Approach

#### De-Cycling

In the first step, we remove cycles from the directed graph, transforming it into a tree, by duplicating the node that completes a cycle, similar to the approach of Mäkinen et al. [46]. If the duplicated node has children, we attach all children to one instance, while the other instance remains a leaf. Figure 3(a) shows a treelike graph with one cycle, Figure 3(b) shows the resulting tree, where node 7 is duplicated. Although this duplication strategy works for general directed graphs, it is most useful for directed graphs with a defined root and few cycles, since in these cases most of the topology is retained, and the number of additional nodes is negligible with respect to scalability.

**Fig. 3.**
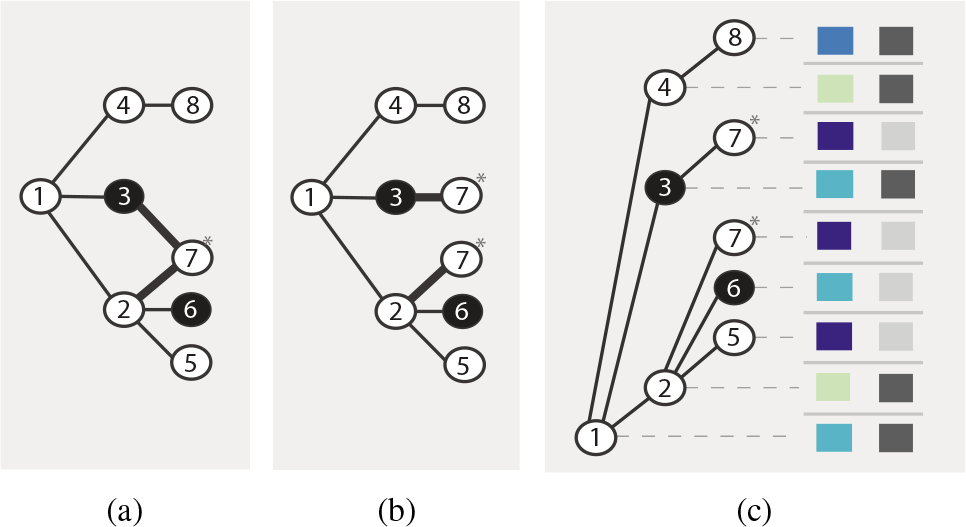
Decycling and linearization. (a) A directed, rooted graph with one cycle ending in node 7. (b) We remove the cycle by duplicating the last node in the cycle (node 7). (c) The tree is linearized so that each node is assigned a distinct row. Leaves are rendered above their parents. This row-based, linear layout enables an unambiguous, position-based association with a table visualizing attributes.

#### Linearization

In most tree layouts [1], associating the nodes with rows in a table by position is impossible. The tree in Figure 3(b) is compact, yet would require, for example, curved links to associate the nodes with a table row. To make this association between nodes and rows of a table intuitive, we use a linearization strategy that assigns every node a distinct vertical position (i.e., a “row”). The position of the node alone thus unambiguously associates the node with a row in a table (see Figure 3(c)). Note that although we assume a left-to-right tree layout here, a top-to-bottom layout would work equally well for associating a tree with table columns.

Linearized tree layouts are based on tree traversal strategies. Although various strategies, such as breadth-first (level-order) or in-order depth-first-search, are possible, we found that a preorder depth-first-search works well for our purposes, since it results in a crossing-free layout and keeps leaves in subsequent rows.

Following the in-order strategy, we recursively place the descendants of a given node directly above their parents. Note that a top-down strategy would also be possible. We assume that an order of leaves can be defined, e.g., based on the attributes. If not, using a random order is possible.

Figure 3(c) illustrates the results of this algorithm when applied to the tree in Figure 3(b) and also shows how to easily associate a table with the tree. Note that the duplicate node also is duplicated in the associated table.

### 5.2 Aggregation

Although linearizing the tree allows for a direct, position-based association of the nodes and their attributes, the resulting layout uses more space than a compact layout. However, due to their hierarchical structure, trees are well suited for aggregation. Degree-of-interest (DOI) functions [20] have been widely applied to trees. In our design, we use the generalized idea of degree-of-interest functions by Furnas [20], [21].

We let analysts define a degree-of-interest function based on the attributes of the nodes, which we call the **phenotype of interest (POI)**. Nodes that have the POI are referred to as **nodes of interest**. In contrast to the original formulation of a degree of interest, our POI function is binary (i.e., a node is either of interest or not) and does not consider a distance to a selected node. An example for a POI is “committed suicide”, which marks all nodes representing individuals who committed suicide to be of interest, or “has a maximum BMI of higher than 30”, which would consider all obese individuals to be of interest. POIs that are a compound of multiple attributes (high BMI and suicide) are possible.

Based on this degree-of-interest function, we introduce two approaches to aggregation that vary in how they trade off compactness and preservation of the attributes of the nodes: (1) attributepreserving aggregation, and (2) attribute-hiding aggregation. These aggregation approaches can, of course, be applied not only to the whole tree, but also to selected subtrees, or to both.

#### Attribute-Preserving Aggregation

Here we introduce an aggregation strategy for linearized layouts that preserves both the structure of the tree and the attributes of all the nodes. Nodes of interest are assigned a row of their own, whereas other nodes are aggregated into a single row. Figure 4(a) shows an example of this strategy applied to the tree shown in Figure 3(c). This layout emphasizes the nodes of interest, while preserving both the structure of the graph and the attributes of the other nodes.

**Fig. 4.**
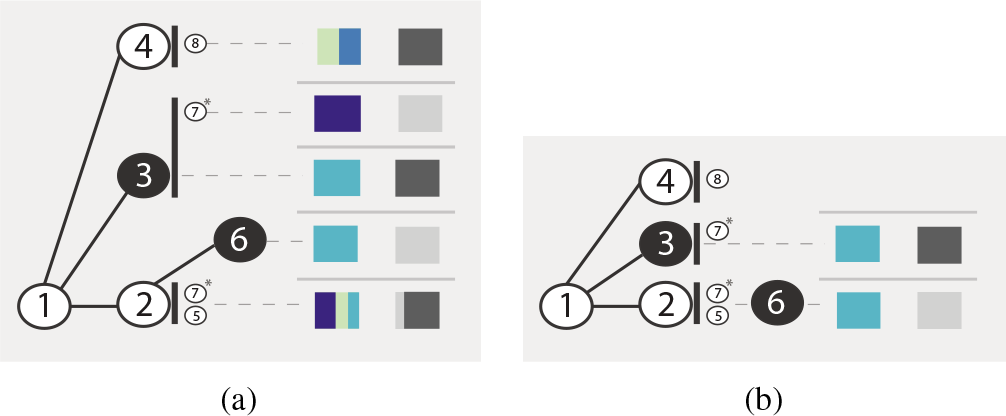
Aggregation approaches demonstrated using the tree in Figure 3(c). A filled-in circle indicates a node of interest. (a) Attribute-preserving aggregation. Each node of interest (shown in black) is in a separate row. Branches without nodes of interest are aggregated into one row, yet all attributes are preserved in the aggregate representations in the table. Notice how the two children of node 2 who are not affected are shown using an implicit encoding, which we refer to as a “kid grid”. (b) Attribute-hiding aggregation. The branches leading to nodes of interest are hidden behind them. Only nodes of interest and branches with no nodes of interest have a row of their own. Only the nodes of interest are represented in the table.

Our algorithm recursively follows a (sub)tree down a branch by assigning a new row to each inner branch. Inner branches are branches that do not end in a leaf after the first edge, i.e., an edge that directly connects to a leaf is not an inner branch. If no node of interest is encountered, the algorithm continues to the leaves, placing all nodes of the branch in the same row. Multiple leaves that are not of interest are placed in a *kid grid*, an implicit encoding of the leaves as small nodes to the right of their parent. These nodes retain all visual encodings (e.g., shape for gender, crossed-out for deceased). We chose this approach for the representation of kid grids over alternative designs such as a numeric labels or bar charts since it is consistent with how individuals are represented in other places in the tree. An example is visible in the bottom branch in Figure 4(a), where nodes 1, 2, 5, and 7 are on the same row, and the leaves (5, 7) are in a kid grid. If a node of interest is encountered, we distinguish two cases. If the node of interest has children that are leaves and that are not nodes of interest themselves, they are added to a kid grid, which is placed in the next row (see node of interest 3,) and its descendant (node 7), which is placed in a kid grid in Figure 4(a)). If the node has children that are inner nodes, the algorithm is applied recursively. The result of this algorithm is a layout that has N rows, where N is the sum of:

- the number of nodes of interest,
- the number of inner branches that do not end in nodes of interest (case for node 4 in Figure 4(a)),
- the number of nodes of interest that have children that are leaves (case for the child of node 3 in Figure 4(a))

The result in the associated table visualization is that each node of interest has a separate row, and the aggregated branches are represented in aggregated rows. In practice, we use visual encodings for aggregates and individual rows that can faithfully represent the data but are also comparable. For details on the table design, see Section 6.2.

#### Attribute-Hiding Aggregation

This form of aggregation also preserves the complete structure of the tree, but it does not preserve attributes of nodes that are not of interest. The result, illustrated in Figure 4(b), is a scalable approach that can be used to address tasks that are concerned only with the attributes of the nodes of interest and their connectivity, but not with the attributes of the other nodes.

The main difference compared to the attribute-preserving aggregation is that nodes of interest are not assigned to a new row when they are encountered. The algorithm again recursively follows a (sub)tree down a branch by assigning a new row to each inner branch. If no nodes of interest are encountered while traversing the branch, the leaves are placed in a kid grid. If a node of interest is encountered, the next step depends on whether it has children that are inner nodes or not. For the node’s children that are leaves, a kid grid is used, but no new row is started. For all other branches, the algorithm is applied recursively.

The resulting layout has M rows, where M is the sum of:

- the number of inner branches,
- the number of nodes of interest that have at least one child that is an inner node.

Here, only nodes of interest and inner branches that do not end in a node of interest are assigned their own row. For consistency, we do not represent branches that do not end in a node of interest in the table.

## 6 LINEAGE DESIGN

Here we describe the design decisions that are specific to the use case of visualizing genealogies and that we realized in the Lineage prototype. To address the tasks of our collaborators, Lineage provides three views, shown in Figure 1: the genealogy graph view, the closely synchronized table view, and a family selection view, which allows analysts to select one or multiple families.

### 6.1 Genealogy Graph

An important difference between genealogical trees and general trees is that nodes have not one but two parents. To address this, we introduce the concept of a couple, indicated by a line connecting the partners (see Figure 5(a)). As is common in genealogical graph layouts, the children of a relationship then connect to the line representing the couple instead of directly to the parents. We also adopt some of the conventions for drawing genealogical graphs: males are drawn as rectangles, females as circles. Deceased individuals are crossed out. In Figure 7(b), for example, the topmost node represents a female who is alive and has the POI, but the other nodes with the POI are deceased. Nodes that have the phenotype of interest are filled in.

**Fig. 5.**
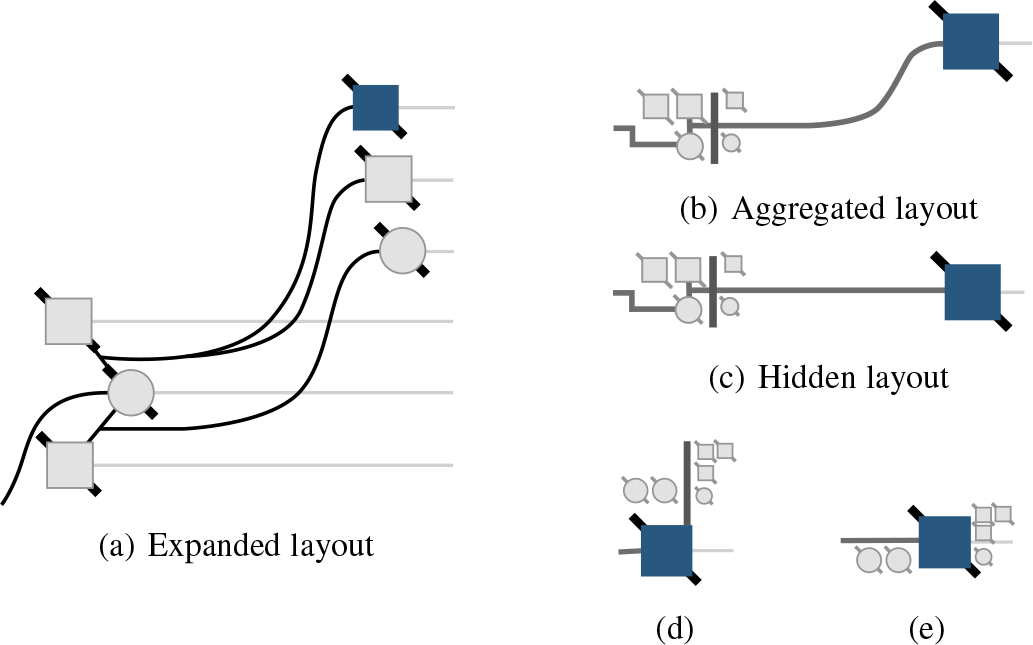
Different aggregation cases. (a-c) A family where one woman has children with two men. One of the children committed suicide. (a) No aggregation: every person is in his or her own row. (b) Attribute aggregation: the suicide case is in its own row; the rest of the family is aggregated. Notice the family grid with two male and one female parents, and one daughter and one son. The second son is not in the kid grid because he is a node of interest. (c) Attribute hiding: the family is hidden behind the suicide case. Only the attributes of the suicide case will be shown in the table. (d-e) A different family, where the node of interest has children, leading to special cases. (d) Attribute aggregation: the spouses and children are moved to their own row. The line connecting spouses spans two rows. (e) Attribute hiding: the spouses are placed to the left of the suicide case, the children to the right.

As discussed in the previous section, the phenotype of interest can be defined dynamically, based on either combining categorical values or brushing a range of a numerical variable. Figure 7 shows the effect of two different POI functions on the same subtree.

The modifications to the layout algorithm to accommodate couples are minor: couples are always placed in consecutive rows to avoid long, vertical parent edges. When one of the spouses has offspring with multiple partners, we place all partners in consecutive rows. In the case of two partners, we place the person with multiple relationships in the center to avoid edge crossings. Figure 5(a), for example, shows a woman who had children with two partners. For more than two spouses, however, or spouses who had children with different partners in alternating order, edge crossings are often unavoidable. Similar to Mäkinen et al. [46], we use arrows to indicate that a node is duplicated and to point toward the duplicate. To resolve any ambiguities, we draw an edge connecting the duplicates when hovering over the arrow (see Figure 6).

**Fig. 6.**
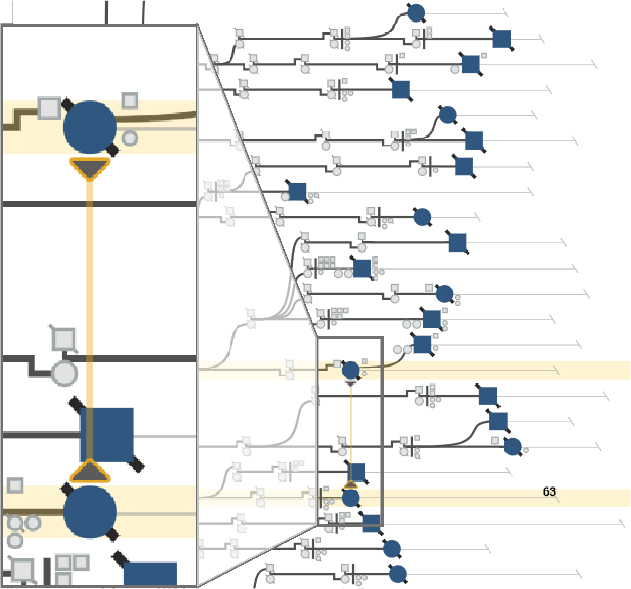
Visual encoding of nodes that were duplicated in the process of removing cycles from the graph. The arrow glyph, which is shown at all times, indicates both the presence of a duplicate and its direction in the graph. Hovering over the arrow draws a line connecting the node to its duplicate and highlights the corresponding rows in the table.

**Fig. 7.**
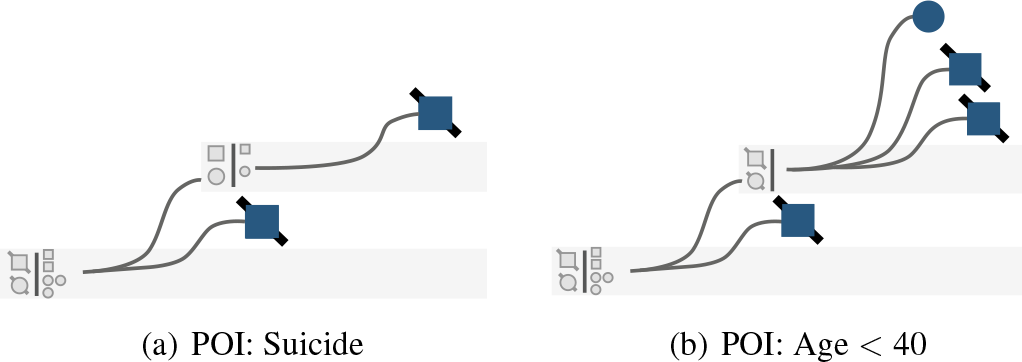
Different POI functions applied to the same aggregated subtree. (a) Suicide as a categorical POI. (b) Age < 40 as a numerical POI.

In contrast to traditional genealogical graphs, we do not lay the nodes out by generation, but use the birth year to position the nodes horizontally [3], as shown in Figure 1. This approach avoids ambiguities about the birth order and encodes a vital attribute directly in the graph. We also use curved splines instead of the traditional orthogonal edge routing, because continuous edges are easier to follow [75].

#### 6.1.1 Aggregation Layouts

With respect to aggregation, the algorithm is extended only by first looking for spouses before descending into a subtree. If both are nodes of interest, each spouse is assigned his or her own row.

We previously introduced the concept of kid grids for aggregated nodes. Indicating hidden nodes using a glyph has been done before for graph layouts, most notably by McGuffin and Balakrishnan [50], who use dots to indicate children in genealogical graphs. Our layout for aggregated genealogies, however, goes beyond a basic indication of existing nodes as they encode both topological information and attributes. First, we extend the notion of a kid grid that encodes children to a family grid that encodes all members of a family. Figure 5 shows multiple examples. A family is separated by a vertical line into parents and children. This vertical line represents the line used to connect spouses in expanded mode. Parents are placed on the left of the line. In addition to the node shape, we also redundantly encode sex by position, placing the nodes representing males on top and the nodes representing females below. In families with multiple partners, we place all partners in the same family grid, so that, for example, a family with a woman who has children with three partners is represented by three squares on top and one circle at the bottom.

Note that aggregation results in some information loss. For families in which individuals have offspring with multiple partners, the exact association between children and parents is lost. Also, the attributes for all aggregated nodes in a row are displayed together in the table, removing the exact association between individuals and their attributes; instead, the distribution of values in that aggregate is emphasized. When hiding is used, the attributes are removed entirely from the table. We found that neither of these drawbacks is a problem since these design decisions align with the analysis tasks outlined earlier: analysts at first are often interested in nodes with the POI. When they want to consider other nodes in detail, they can deaggregate on demand.

It is important to note that we break with the convention of placing nodes based on their birth-year for aggregated families. Instead, we place the whole family based on the birth year of the parent with a blood relationships to the ancestors.

#### 6.1.2 Encoding Attributes in the Graph

Although we address the problem of encoding multiple attributes for nodes using our linearization approach, direct, on-node encoding of a small number of attributes provides the best bridge between attribute-based and topology-based tasks. We already discussed how sex (shape), deceased/alive (crossed out), birth year (horizontal position), and POI (fill) are encoded directly in the graph. To enable our collaborators to view an additional variable in the graph, we introduce a glyph, rendered to the right of the nodes, as shown in Figure 8. When the attribute is categorical, we color-code the glyph; for numerical attributes, we show a small bar. In both cases, the color coding is also used in the table to highlight the relationship (see the matching colors for bipolar disorder in Figures 8(a) and 9). When data is not available for a node, no glyph is shown.

**Fig. 8.**
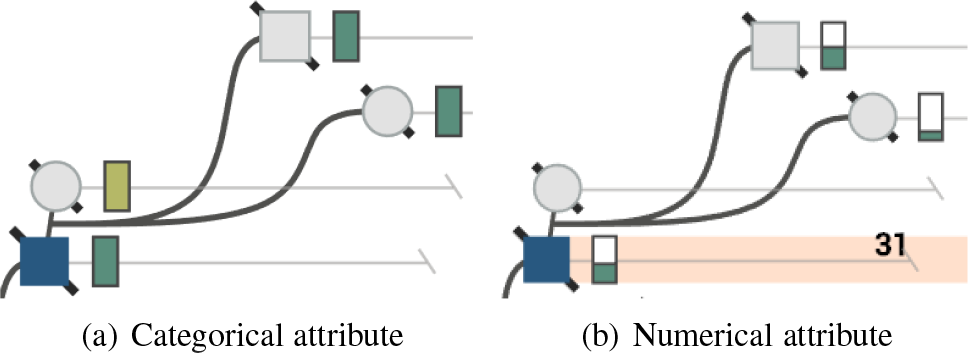
Attributes encoded directly in the graph. Age lines visualize the lifespan of individuals. Age lines for people who are alive continue until the present. Age lines of deceased individuals are terminated at their year of death. We can see that the individual represented by the node of interest died at age 31, and his spouse died shortly thereafter. Selected attributes can be visualized next to the nodes in glyphs (green rectangles). (a) The categorical variable bipolar disorder is encoded by a dark-green color. (b) The numerical variable number of bipolar diagnoses is encoded as a bar chart.

Finally, we also encode the age of individuals directly in the graph by drawing a line from the node, which is placed at the year of birth, to the year of death, or to the current year (see Figure 8). These age lines conveniently encode an important variable in the existing coordinate system. We found that the age lines also help to perceptually connect the nodes to the rows in the table. Since we do not draw age lines for aggregates, we found it necessary to indicate the connection to the table using a light-gray background, as can be seen, for example, in Figure 7.

**Fig. 9.**
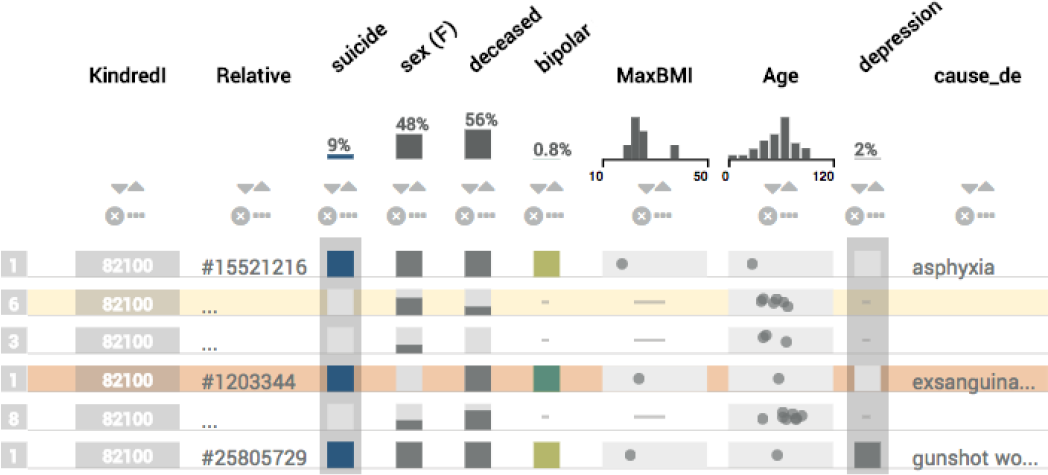
The table view. The first column encodes how many individuals are aggregated in that row. Binary categories are represented as present/absent (e.g., sex). Aggregates of binary variables show the proportions of the variable in stacked bars. Numerical values are encoded using dot plots, which are also used for aggregates. The POI is highlighted using a gray background. The depression column is starred, also indicated by the gray background.

### 6.2 Table Visualization

The attribute table is designed to visualize both rows representing individuals and aggregates representing multiple individuals in the same space. The attributes visualized in the table can be chosen using the *Table Attributes* menu in the tool bar. As shown in Figure 9, we use dot plots to encode numerical data. Combined with transparency and jitter, dot plots can also be used to encode aggregate rows. For categorical values, we distinguish between binary categories, such as deceased or alive, and multivalued categories, such as race. We encode binary categories in a single column, as can be seen for “sex (F)”, where a dark cell corresponds to true and a light cell corresponds to false. For multivalued categories, we use a method commonly employed by Bertin [8], and use one binary column for each category instead of, e.g., using color to encode categories. We represent aggregates of binary or categorical values as stacked bars, which are scaled according to the number of individuals in a category. Text labels and IDs do not have adequate visual representations for groups of elements, so we display an ellipsis (…) for aggregates. Missing values are rendered as a dash to distinguish them from zero or false values. We also provide a column that shows how many people are in a given row.

We avoid color to encode data, so we can employ it to highlight elements of interest, such as to highlight selected rows and to indicate the column that encodes the user-selected phenotype of interest and the primary attribute. In Figure 9, the selected attribute (bipolar) and the POI (suicide) are rendered in color. A menu in the columns allows analysts to set the POI, set an attribute as a primary attribute, and *star* an attribute. Starring an attribute adds it to the family selector table.

These features, in combination with the graph, allow analysts to address the tasks related to analyzing individuals (**T2**) and comparing cases (**T3**).

Finally, we also allow analysts to sort the table based on any column, which enables them to to easily identify clusters of similar items (**T4**). However, sorting by attribute removes the close association with the graph. To partially remedy this problem, we draw slope charts, similar to what is used in LineUp [26], to relate the rows of the table to the rows of the graph. These connection lines work well for a small number of rows, but often result in significant crossings when dealing with many rows. In that case, interactive highlighting helps to trace the lines. Lines that end in an off-screen location are not rendered. Instead, an icon indicates the direction of their corresponding row. Clicking on the icons automatically scrolls to the location of the corresponding row in the table. Figure 10 shows an example of a partially aggregated graph sorted by suicide.

**Fig. 10.**
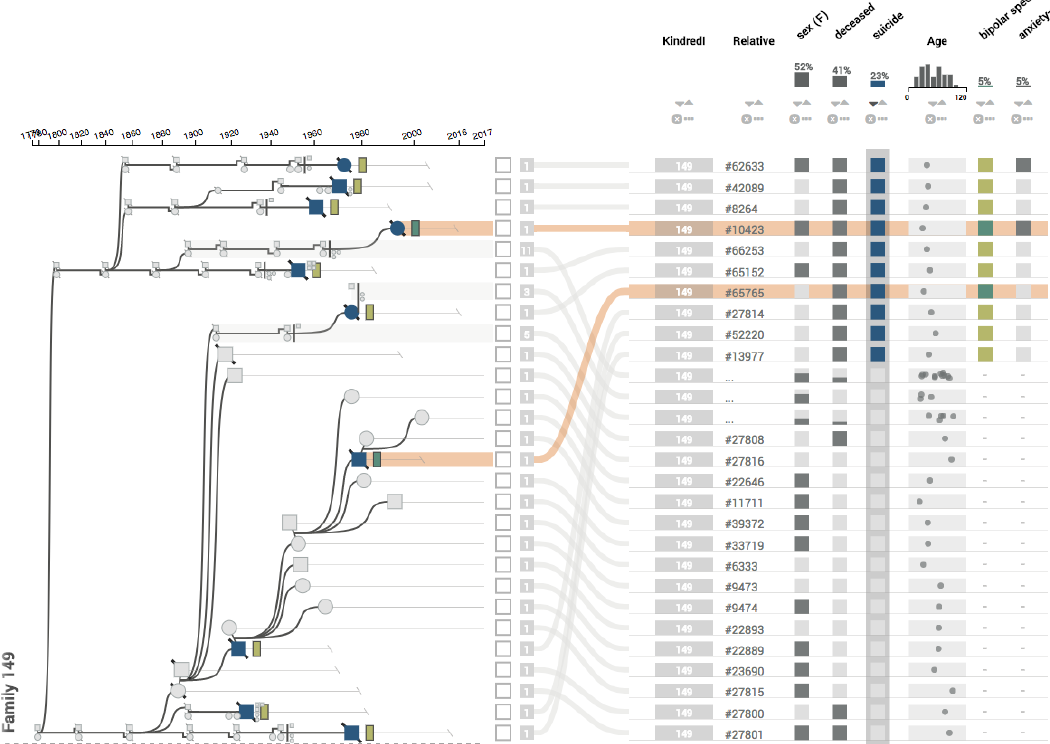
The table is sorted by suicide, which causes the rows in the table to be in a different order than the rows in the graph. The association between the two is retained by the curves connecting them.

### 6.3 Viewing Multiple Families

One important aspect of our collaborators’ workflow is the comparison of multiple families (**T5**). A requirement for comparison of families is the ability to select families **T1**, which is enabled by the family selection view, shown on the left in Figure 1. The family selection view shows statistics about the family, such as its size, the number of people with the currently selected POI, and the number of people who have a starred attribute. Combined with sorting, this feature is useful to identify families with a high incidence of an attribute, for example, to identify families in which bipolar disorder is common. Multiple families are seamlessly integrated into the graph and table views (see Figure 11). To visually separate the families, a dashed line is drawn between them.

**Fig. 11.**
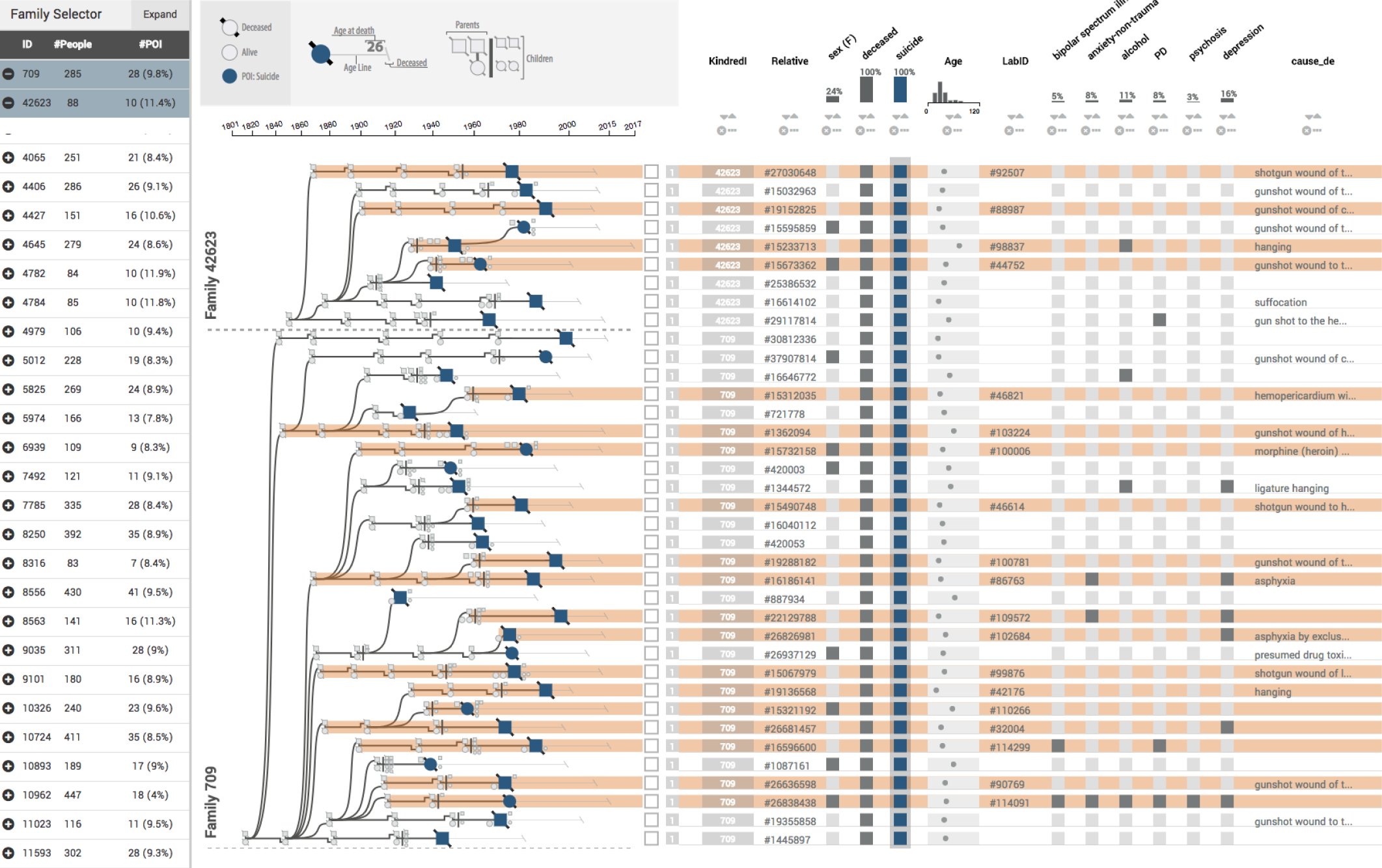
Two families with high familial risk for suicide. Our collaborator uses Lineage to prioritize families for an analysis of shared genomic sequences of suicide cases. Although both families seem promising based on risk alone, Lineage reveals that the few cases with genetic material (indicated by the presence of a *LabID*, highlighted in orange) in Family 42623 are too closely related. Family 709, in contrast, contains 15 cases with genetic material available that are also widely separated. Furthermore, a large cluster of relevant comorbidities (depression, anxiety, alcohol abuse, bipolar disorder, personality disorder) indicates a likely genetic component of suicide in this family.

## 7 IMPLEMENTATION AND PREPROCESSING

Lineage is open source, is implemented in TypeScript as a Caleydo Phovea client/server application [25], and uses D3 for rendering. The server component is based on Flask and is provided as a Docker container for easy deployment. A prototype of Lineage is available at https://lineage.caleydoapp.org. The source code is available at https://github.com/caleydo/lineage.

The prototype made available publicly contains 10 selected and anonymized families from the suicide study based on data from the Utah Population Database. The anonymization method and the selection of families were approved by the Utah Resource for Genetic and Epidemiologic Research (RGE). The anonymization process involves randomizing the sex of individuals and the birth and death years and randomly deleting individuals, in addition to omitting attributes that could be identifiable. Hence, we do not recommend making clinical inferences based on the data provided.

## 8 CASE STUDIES

We present two forms of validation for Lineage: case studies, a method employed widely to demonstrate the fitness for use of visualization design studies [37], [64], and informal usability testing and analyst feedback, which is described in the next section.

The case studies outlined below were conducted with Dr. Hilary Coon, a psychiatry researcher and principal investigator studying the genetic and environmental factors in suicide. Dr. Coon, who is a coauthor of this paper, also participated in the analyst feedback sessions described in Section 9, and contributed to the design and development of Lineage. She was familiar with the interface from the earlier analyst feedback sessions as well as through demonstrations during the design phase. For the case study, we deployed Lineage on a secure, password protected and HIPPA compliant server instance on Amazon Cloud Services, which allowed her to access the tool from her personal computer, in her own work environment, and at a time that was convenient.

For the case studies, Dr. Coon used the full dataset described in Section 2 and performed analysis tasks representative of her research. She documented her analysis process with notes and screenshots, which she shared with the team. Sections 8.2 and 8.3 were written by Dr. Coon and reflect her detailed accounts of the analysis workflow. These case studies were subsequently used in a research grant proposal.

These case studies demonstrate how Lineage allows our collaborators to complete several tasks, including (1) selecting families most likely to point to genetic factors of suicide, and determining additional defining characteristics of cases in these families for familial analysis (**T1**); (2) visualizing aspects of familial cases contributing to genomic shared regions with genome-wide significant evidence (**T3**); (3) searching for supporting evidence by visualizing additional families with evidence of familial sharing in the same genomic region (**T5**); and (4) prioritizing new families for additional analysis based on visualization of risk and clustering of defining characteristics (**T4**).

### 8.1 Prioritizing Families for Analysis

Dr. Coon uses the suicide dataset described in Section 2 for her analysis. Using established familial relative risk methods [31], she ascertains a large number (>200) of extended multigeneration high-risk families for analysis [14]. Limited time for analysis and resources requires her to prioritize these families based on data visualized in Lineage. Our collaborator’s goal is to select a promising family for the computationally expensive analysis of Shared Genomic Segments (SGS) [74]. SGS investigates the significance of genomic segments shared between distantly related affected cases. If an observed shared segment of the genome is significantly longer than expected by chance, then inherited sharing is implied. A greater distance separating cases (i.e., path-distance in the genealogy graph) translates to the increased statistical power of the method: chance inherited sharing in distant relatives is improbable. Finding cases with shared sequences that also have a shared phenotype (suicide, plus potentially other comorbidities) allows Dr. Coon to reason that the sequence is a factor in these phenotypes.

Previous analyses exploring the statistical power of SGS using simulated high-risk families indicated that power is determined by familial distance (path length) between cases with genomic data for analysis, indexed by counting the total number of generations separating the cases (meioses). In this study, if families had at least 15 meioses between cases, then 3-10 families were sufficient to gain excellent power (>80%) to see at least one true positive within any given family [35]. For all scenarios considered, genome-wide association studies (i.e., studies that do not consider familiarity) would have negligible power to detect the simulated variants. This study therefore dictated that extended high-risk families should be selected with the highest familial relative risk, and with the largest number of cases with genotyping available for analysis through interrogating numbers of cases with DNA, and familial distance of these cases from one another. Familial distance is visualized in the graph view in Lineage and serves as a useful predictor of the statistical significance of shared regions among cases. However, additional data about the cases shown in the table view of Lineage is useful in later prioritization of genes in regions with significant evidence of sharing. These attributes include gender, young age at death, and clustering of comorbidities.

Figure 11 shows two families of high interest (709, 42623), based on significant familial risk ratios (p<0.0001). These familial risk ratios are calculated with a separate tool. Significant families are selected by ID in Lineage. Both families have more than three cases available for analysis, as indicated by the *LabID* attribute. However, 42634 was not chosen for analysis given that cases are not sufficiently separated (of the four cases with a LabID, two have a parent-child relationship, for instance), and little cooccurring diagnostic information is apparent in this family. Lineage reveals that family 709, in contrast, suggests clustering of multiple conditions, and has 15 suicide cases with genetic material available that are also adequately separated (**T4**).

Our collaborator hence selected family 709 for the computationally intensive analysis of shared genomic segments. Creation of files for this analysis was facilitated by exporting the LabIDs from the family, which serve as the input “proband list”.

### 8.2 >Identifying Comorbidities of Cases Contributing to Significant Familial Genomic Sharing

Analyses of selected families to date have produced significant evidence of genomic sharing. An example is a shared region in family 601627 with genome-wide significance (p=1.94E-10). A single gene in this region, *neurexin 1 (NRXN1)*, has evidence from the published literature of involvement in psychiatric risk [48] and inconclusive evidence for risk of suicide [54]. The shared region in family 601627 is shared by six cases, shown in Figure S4 in the supplemental material. Dr. Coon used Lineage to interactively explore demographic attributes and clustering of clinical cooccurring conditions (**T3**). Cases contributing to sharing had young age at death, ranging from age 17 to 39; average age at death in the research cohort is 40. The clustering of co-occurring depression is not completely unexpected, as approximately half of the suicide cases show evidence of depression. However, the multiple cases with personality disorders (PD) are more unexpected. These disorders include less commonly used diagnoses such as antisocial personality, borderline personality, conduct disorder, and obsessive personality.

This association was previously unknown to Dr. Coon, and would have been very difficult to identify given the number of cases in the family (831) and the number of case attributes. Knowledge of clustering of attributes in these cases with evidence for this particular genetic risk will direct our collaborator in her selection of additional families for resource-intensive replication studies. The attributes will also guide selection of other nonfamilial cases from their much larger cohort and of external cohorts for further replication. If replication is achieved, knowledge of case attributes could also drive the design of additional targeted studies in specific high-risk subgroups.

Based on these discoveries, our collaborator then extended her search for other families associated with shared regions that are associated with the gene *NRXN1* (for details, see Section 2 in the supplementary material). She found two more families associated with the gene *NRXN1* and consistent demographic attributes and comorbidities (Figure S5 in the supplement). Across all three families, the occurrence of PD in the cases supporting the genomic regions was 8/19=42%. Once the three families had been visualized, another feature was observed (**T5**). Of the 19 cases across all families, 6 were female (31.6%). The percentage of female cases in the overall cohort was 20.8%. The occurrence of this trend for this important demographic attribute will help with selection of additional families to continue to follow up evidence for this and related genes.

### 8.3 Prioritizing Additional Families

Our collaborator has found Lineage to be particularly helpful in prioritizing additional families to follow up initial results based on clustering of important attributes. For the case study above, the important attributes were young age at death, presence of PD, enrichment for female cases, and possibly presence of depression. Within these selection attributes, families had to also meet minimum requirements of number and separation of cases with available data, as discussed in the first case study (**T1**).

Our collaborator started by sorting the families by the occurrence rate of personality disorders (PD) within Lineage, which resulted in relatively few families with a concentration of cases with these disorders. She looked not only at overall percentage of suicide cases in the families with PD, but also at numbers of cases with DNA for analysis; even if a family has overall high enrichment of this diagnosis, if only one or two cases have genetic data, the family will be of little interest. Given this initial filter, three families were prioritized at the top (**T1**): families 540781 (7 PD cases total, 3 with DNA), 10724 (6 PD cases total, 4 with DNA), and 565350 (5 PD cases total, 3 with DNA). Figure S6 in the supplement shows the attribute table for these families. As Dr. Coon was performing this initial task, she noticed that a high number of the PD suicide cases were also women, even in families not at the top of this follow-up priority list (**T4**). Although the association between these attributes was present in the family described in the previous case study (6 of the 19 PD cases supporting sharing in the initial three families were female), this signal was even stronger in these families identified based on the phenotype characterized. In families 540781, 10724, and 565350, the proportions of female PD cases were 4/7, 2/6, and 5/5, respectively. Taking all families together, the proportion of female cases among those with PD was 11/18 = 61%. Given that female suicide makes up only about 20% of cases overall, this observed pattern is striking. Our collaborator is now interested specifically in this association as it may relate to *NRXN1* genetic risk, but also more generally in the association of these attributes and how this may relate to other phenotypic and genetic aspects of the research sample.

It became apparent to her that the initial prioritization already fulfilled another criterion, enrichment for female suicide, likely due to the association between this characteristic and the occurrence of PD. Note that this enrichment for female suicide is apparent only when also considering the PD attribute; taken as a whole, these families are not significantly enriched for female suicide.

Another attribute of interest was age at death. Taking all age at death for cases with DNA in families 540781, 10724, and 565350, Dr. Coon found averages of 32.86 (sd=12.48), 43.42 (sd=13.98), and 26.33 (sd=9.06), respectively (**T3**). This factor suggests that families 565350 and 540781 may be most interesting for computationally intensive follow-up of the initial findings, and will help her and her team makes decisions regarding which cases to select for expensive molecular sequencing.

Overall, the case study revealed several ways in which Lineage aided in the analysis process, as detailed above, as well as a few limitations of the tool. One such drawback is the need to identify families of interest prior to using Lineage in order to select relevant families for comparison within the tool. Another limitation is the need to manually count selected individuals in order to calculate the percentage of a certain family represented by a selection. These limitations are addressed in further detail in Section 11.

## 9 ANALYST FEEDBACK

In addition to the case studies, we ran an informal feedback session with two faculty members, one research scientist, and one PhD student. With the exception of one faculty member, who is also a co-author, the participants did not contribute to the design and development of Lineage, except for the requirement analysis as described in Section 3.

After an introduction to Lineage, the participants were asked to use the tool with their own data and articulate their thought process and observations according to the think-aloud protocol, followed by a brief interview. The sessions took between 90 minutes and two hours.

The feedback we received was overwhelmingly positive, including statements such as *“This is going to completely change how we do things*”. One analyst noted that Lineage will allow him to properly use visualization for the exploration of genealogies for the first time because their current tools are not suitable for discovery, since they can only effectively visualize one or two attributes at the same time, and the tools are essentially static and difficult to use.

The analysts consistently noted that the integration of attributes and family structure is critical for them to make decisions about where to follow up with subsequent analysis, making comments such as “*I think it’s really helpful to see the attributes next to the graph. It really helps to pinpoint the important cases*”.

We asked the analysts about their opinions on attribute-preserving aggregation and how it compared to attribute-hiding aggregation. They commented that attribute-preserving aggregation is not particularly useful for their suicide dataset due to the sparse attributes of the nonaffected individuals, but that they can imagine it might be very useful when applied to their autism dataset, which contains more data on family members. One analyst gave the example that he would be interested to see autism spectrum scores aggregated for a whole family.

The analysts also stated that they believe that Lineage graphs are appropriate for presentations in publications and presentations, as the visual encodings are easy to explain. They asked for some features in support of presentations, such as the ability to hide irrelevant branches or nodes of the graph, or to redefine the founder to clean up the genealogy. Finally, we also asked for other features that they wished the tool had. The answers to that were mostly regarding data, i.e., to load more data into the tool and to provide export capabilities for a subsequent statistical analysis. Also, a search features for individuals and families was mentioned by multiple participants.

## 10 DISCUSSION

Although details of our design study and our implementation, such as how we display parents and family grids, are specific to genealogies, we argue that our linearization and attribute-driven aggregation approach can be applied broadly when analyzing multivariate trees or tree-like graphs, such as phylogenies or file directories. The species and their relationships depicted in phylogenies, for example, are associated with vast numbers of attributes capturing traits (is flightless, has tail, color, etc.), and judging which attribute is inherited at which point in the tree of life is crucial for understanding the process of evolution. Answering these questions is important in a basic science context as well as in a human health context. An example for the latter is the study of the development of viruses such as influenza, Zika, or HIV [57]. Our approach also has the potential to be combined with more generic graph-to tree extraction approaches, as discussed by Lee et al. [40]. Using their method, arbitrary multivariate graphs could be converted into trees and explored in Lineage.

Lineage as a clinical genealogy visualization tool specifically can be applied to study other diseases with a major impact on human health. We have already deployed Lineage with an autism dataset (see supplemental Figure S3), which has characteristics that emphasize the importance of attribute visualization of nonleaf nodes. In this dataset, attributes are also available for parents and other relatives of autism cases.

We also argue that our strategy of combining explicit node-link layouts with the implicit layout of the family grids is transferable to other application scenarios.

Our described linearization approach makes the association between nodes and attributes obvious and enables a tight integration of attribute-based and topology-based graph analysis tasks. Both aggregation methods described serve to reduce the space usage of the linearized tree while preserving the topology and the desired level of information about the attributes. The aggregation is based on two principles: assigning nodes to be aggregated to the same row and combining the explicit node-link layout with the implicit encoding for aggregated nodes and their leaves (family grids).

The Lineage genealogy visualization tool can be broadly used with other genealogical datasets, e.g., to study autism, diabetes, or cancer. Many groups at the University of Utah make use of the Utah Population Database, and we have already established contact with other potential collaborators who are in need of a clinical genealogy visualization tool. Some of these datasets also have detailed attributes for nonaffected cases, which will make our attribute-preserving aggregation approach more valuable. Although our data is unique with respect to its scope, detailed genealogical datasets are becoming more common because they have shown immense potential for population genetics [47]. We believe that our approach could also be adapted to datasets containing many small families (siblings, parents, grandparents of affected individuals) since they are commonly collected to study the genetic disease of one family member.

## 10.1 Scalability

In contrast to other tools, such as the DOITree [27], our aggregation approach preserves all the structure of the tree, which is suitable for trees with hundreds of nodes, but not for trees with tens of thousands of nodes or more. To scale to larger trees, our algorithms could be combined with hiding parts of the tree. Also, although our algorithms work for any tree and any phenotype of interest, they are most efficient if the number of nodes of interest is small compared to the number of nodes in total. A common phenotype of interest for our collaborators is suicide, and the typical genealogies they study contain between 5-15% suicide cases. For these conditions, we found the resulting layouts to be compact and useful.

We found Lineage to scale well to families with about 1500 individuals, which covers most families in our collaborators’ dataset (547 of 550 families have fewer than 1000 individuals). We also experimented with the largest families in our dataset, which contain about 2500 individuals. For these families, we observed several seconds of wait time until the decycling and the layout were computed. We anticipate addressing these performance limitations through precomputing and caching initial layouts.

In terms of the scalability of the visual encodings, we argue that Lineage produces a more readable layout in less space than Progeny, the tool that is currently used by our collaborators for displaying genealogies. Note that Progeny has only very limited capabilities for showing attributes by encoding attributes directly on the nodes and displaying text underneath nodes, and attributes cannot be dynamically selected or manipulated. For a comparison between Progeny and Lineage, please refer to the supplementary document. When using suicide as a POI (the most common use case) and when using attribute-hiding aggregation, a family with about 400 individuals fits onto a single screen without scrolling (see Supplementary Figure S2). Larger families, attribute-preserving aggregation, or no aggregation more commonly require scrolling.

The number of attributes that can be displayed for each individual is limited by the horizontal screen size. On a large, 2560×1600 pixel display, about 20-40 dimensions can be shown, depending on the type (text and numerical columns need more space than binary categorical, for example). We found that this number typically exceeds the number of attributes our collaborators would like to study simultaneously.

## 11 CONCLUSION AND FUTURE WORK

In this paper, we introduced a novel approach for visualizing multivariate trees and tree-like graphs using a linearization approach. We demonstrate the usefulness of our approach by realizing it in the Lineage system, which is designed for the visualization of genealogies in a clinical context. Using Lineage, our collaborators are now able to efficiently explore the structure of large families and even multiple families at the same time, in addition to analyzing dozens of attributes for the individuals in these families. They can use Lineage to identify phenotypes of interest that appear in multiple families, and then use this knowledge to inform and narrow down their search for genetic variants.

Lineage in its current form is already useful to our collaborators, but there are many directions in which it could be extended. Specifically, we currently deal with only a selected subset of the 3000 dimensions that are available for each of our cases. We plan to develop integrated visual and analytical methods to select dimensions of interest for any given subset of patients. For example, the system could identify that for a given family, PTSD is a common comorbidity and suggest that the analyst add PTSD to the table. Such an approach will be especially important when we start to integrate the detailed genetic data that is available for many of these cases.

The case studies and the feedback session also revealed areas for future work. First, it is desirable to integrate search and filter functionality, so that analysts can quickly identify families of interest based on attribute data. In our current implementation, participants used an external spreadsheet with statistical information about the families, and combined it with the browser search feature to find families of interest. A second aspect is a panel that displays basic information about a selection, such as the number of selected individuals, as well as the percentage of the total family selected. In our case study, these numbers were achieved by manual counting.

Finally, we are exploring other application areas for Lineage. We are currently discussing Lineage with evolutionary biologists, who are excited about the potential of our multivariate tree visualization method for trait analysis in large phylogenetic trees.

## ACKNOWLEDGMENTS

We thank Asmaa Aljuhani and Annie Cherkaev for their contributions. We also thank our collaborators and the Visualization Design Lab at the University of Utah for the feedback, and the Caleydo team for their technical support. This work was supported in part by the US National Institutes of Health (U01 CA198935, R00 HG007583, R01MH099134) and the DoD – Office of Economic Adjustment (OEA), ST1605-16-01. We thank the Pedigree and Population Resource of the Huntsman Cancer Institute, University of Utah (funded in part by the Huntsman Cancer Foundation) for its role in the ongoing collection, maintenance, and support of the Utah Population Database (UPDB). We also acknowledge support for the UPDB through grant P30 CA2014 from the National Cancer Institute, and from the University of Utah’s Program in Personalized Health and Center for Clinical and Translational Science.

## Footnotes

1 https://healthcare.utah.edu/huntsmancancerinstitute/research/updb/

2 The terms genealogy and pedigree can be used interchangeably in this context. However, for simplicity, we will always use genealogy

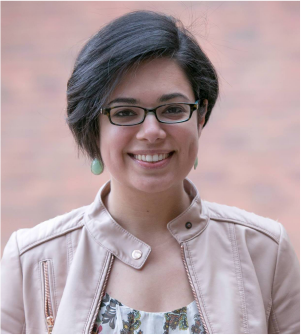

**Carolina Nobre** is a PhD student at the Scientific Computing and Imaging Institute and the School of Computing at the University of Utah. Carolina works under the supervision of Alexander Lex in the Visualization Design Lab (VDL) and is interested in data visualization, particularly related to large multivariate networks.

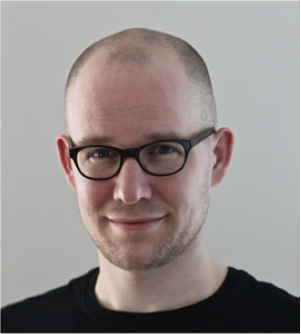

**Nils Gehlenborg** is an Assistant Professor in the Department of Biomedical Informatics at Harvard Medical School. Nils obtained his PhD from the University of Cambridge and the European Bioinformatics Institute (EMBL-EBI). He is interested in visualization of biomedical data, in particular with applications in genomics, epigenomics, and cancer biology.

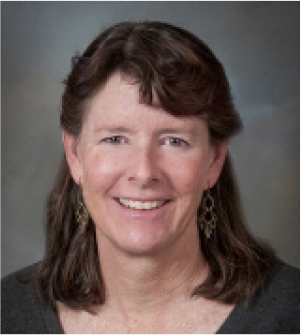

**Hilary Coon** is a Research Professor of Psychiatry at the Department of Psychiatry at the University of Utah. She received her PhD in Psychology from the University of Colorado and Institute for Behavioral Genetics, Boulder, Colorado in 1991. Hilary’s primary research interests include finding genetic risk factors that lead to susceptibility to suicide, and also autism genetic risk. Her interests also include the development and application of statistical methods to genetic and phenotypic data, and genetics of other complex disorders, including cardiovascular disease, obesity, and lung disorders.

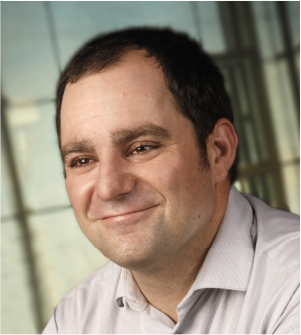

**Alexander Lex** is an Assistant Professor of Computer Science at the Scientific Computing and Imaging Institute and the School of Computing at the University of Utah. Before joining Utah he was a lecturer and a post-doctoral visualization researcher at Harvard University. He received his PhD from the Graz University of Technology in 2012. His primary research interests are data visualization, especially applied to molecular biology, and human computer interaction.

## References

[1] R. Adrian. Tree Drawing Algorithms. In Handbook of Graph Drawing and Visualization, pages 155–192. CRC Press, 2013.

[2] A. V. Bakian, R. S. Huber, H. Coon, D. Gray, P. Wilson, W. M. McMahon, and P. F. Renshaw. Acute Air Pollution Exposure and Risk of Suicide Completion. American Journal of Epidemiology, 181(5):295–303, 2015.

[3] R. Ball. Visualizing genealogy through a family-centric perspective. Information Visualization, 16(1):74–89, 2017.

[4] J. C. Barrett,B. Fry, J. Maller, and M. J. Daly. Haploview: Analysis and visualization of LD and haplotype maps. Bioinformatics, 21(2):263–265, 2005.

[5] A. Barsky,T. Munzner, J. Gardy, and R. Kincaid. Cerebral: Visualizing Multiple Experimental Conditions on a Graph with Biological Context. IEEE Transactions on Visualization and Computer Graphics (InfoVis ’08), 14(6):1253–1260, 2008.

[6] R. L. Bennett, K. S. French, R. G. Resta, and D. L. Doyle. Standardized Human Pedigree Nomenclature: Update and Assessment of the Recommendations of the National Society of Genetic Counselors. Journal of Genetic Counseling,17(5):424–433, 2008.

[7] R. L. Bennett, K. A. Steinhaus, S. B. Uhrich,C. K. O’Sullivan, R. G. Resta, D. Lochner-Doyle, D. S. Markel, V. Vincent, and J. Hamanishi. Recommendations for standardized human pedigree nomenclature. Pedigree Standardization Task Force of the National Society of Genetic Counselors. American Journal of Human Genetics,56(3):745–752, 1995.

[8] J. Bertin. La Graphique et Le Traitement Graphique de l’information. Nouvelle bibliotheque scientifique. Flammarion., 1975.

[9] A. Bezerianos, F. Chevalier, P. Dragicevic, N. Elmqvist, and J. D. Fekete. GraphDice: A System for Exploring Multivariate Social Networks. Computer Graphics Forum (EuroVis ’10), 29(3):863–872, 2010.

[10] A. Bezerianos, P. Dragicevic, J. Fekete, J. Bae, and B. Watson. Genea Quilts: A System for Exploring Large Genealogies. IEEE Transactions on Visualization and Computer Graphics,16(6):1073–1081, 2010.

[11] B. Cannon, M. Hiremath, C. Jorcyk, and A. Joshi. CoVE: A Colony Visualization System for Animal Pedigrees. In Proceedings of the 7th International Symposium on Visual Information Communication and Interaction, VINCI ’14, pages 9:9–9:18. ACM, 2014.

[12] S. K. Card and D. Nation. Degree-of-Interest Trees: A Component of an Attention-Reactive User Interface. In Proceedings of the Working Conference on Advanced Visual Interfaces, AVI ’02, pages 231–245, New York, NY, USA, 2002. ACM.

[13] W. S. Cleveland and R. McGill. Graphical Perception: Theory, Experimentation, and Application to the Development of Graphical Methods. Journal of the American Statistical Association, 79(387):531–554, 1984.

[14] H. Coon, T. M. Darlington, W. B. Callor, E. Ferris, A. Fraser, Z. Yu, N. William, S. C. Das, S. E. Crowell, M. Puzia, M. Klein, A. Docherty, L. Jerominski, D. S. Cannon, K. R. Smith, B. Keeshin, A. V. Bakian, E. Christensen, N. J. Camp, and D. Gray. Identification of genome-wide significant shared genomic segments in large extended Utah families at high risk for completed suicide. bioRxiv, page 195644, 2017.

[15] G. M. Draper andR. F. Riesenfeld. Interactive fan charts: A space-saving technique for genealogical graph exploration. In Proceedings of the 8th Annual Workshop on Technology for Family History and Genealogical Research (FHTW 2008), 2008.

[16] M. B. Eisen, P. T. Spellman, P. O. Brown, and D. Botstein. Cluster analysis and display of genome-wide expression patterns. Proceedings of the National Academy of Sciences USA,95(25):14863–14868, 1998.

[17] D. Engel, R. Rosenbaum, B. Hamann, and H. Hagen. Structural Decomposition Trees. Computer Graphics Forum,30(3):921–930, 2011.

[18] S. Fu, H. Dong, W. Cui, J. Zhao, and H. Qu. How Do Ancestral Traits Shape Family Trees Over Generations? IEEE Transactions on Visualization and Computer Graphics, 24(1):205–214, 2018.

[19] C. Fuchsberger, S. Miksch, L. Forer, and C. Pattaro. Analyzing Populations with Visual and Analytical Methods to Identify Family Clustered Diseases. Proceedings of the 12th World Congress on Health (Medical) Informatics; Building Sustainable Health Systems, page 2243, 2007.

[20] G. W. Furnas. Generalized fisheye views. In Proceedings of the SIGCHI Conference on Human Factors in Computing Systems(CHI ’86), pages 16–23. ACM, 1986.

[21] G. W. Furnas. A Fisheye Follow-up: Further Reflections on Focus + Context. In Proceedings of the SIGCHI Conference on Human Factors in Computing Systems, CHI ’06, pages 999–1008. ACM, 2006.

[22] N. Gehlenborg,S. I. O’Donoghue, N. S. Baliga, A. Goesmann, M. A. Hibbs, H. Kitano, O. Kohlbacher, H. Neuweger, R. Schneider,D. Tenenbaum, and A.-C. Gavin. Visualization of omics data for systems biology. Nature Methods, 7(3):56–68, 2010.

[23] M. Glueck, A. Gvozdik, F. Chevalier, A. Khan, M. Brudno, and D. Wigdor. PhenoStacks: Cross-Sectional Cohort Phenotype Comparison Visualizations. IEEE Transactions on Visualization and Computer Graphics (VAST ’16), 23(1):191–200, 2017.

[24] S. Goodwin, J. Dykes, S. Jones, I. Dillingham, G. Dove, A. Duffy, A. Kachkaev, A. Slingsby, and J. Wood. Creative User-Centered Visualization Design for Energy Analysts and Modelers. IEEE Transactions on Visualization and Computer Graphics,19(12):2516–2525, 2013.

[25] S. Gratzl, N. Gehlenborg, A. Lex, H. Strobelt, C. Partl, and M. Streit. Caleydo Web: An Integrated Visual Analysis Platform for Biomedical Data. In Poster Compendium of the IEEE Conference on Information Visualization (InfoVis ’15). IEEE, 2015.

[26] S. Gratzl, A. Lex, N. Gehlenborg, H. Pfister, and M. Streit. LineUp: Visual Analysis of Multi-Attribute Rankings. IEEE Transactions on Visualization and Computer Graphics (InfoVis ’13), 19(12):2277–2286, 2013.

[27] J. Heer and S. K. Card. DOITrees Revisited: Scalable, Space-constrained Visualization of Hierarchical Data. In Proceedings of the Working Conference on Advanced Visual Interfaces, AVI ’04, pages 421–424, New York, NY, USA, 2004. ACM.

[28] B. Johnson and B. Shneiderman. Tree-maps: A space-filling approach to the visualization of hierarchical information structures. In Proceedings of the IEEE Conference on Visualization (Vis ’91), pages 284–291, 1991.

[29] I. Jusufi, Y. Dingjie, and A. Kerren. The Network Lens: Interactive Exploration of Multivariate Networks Using Visual Filtering. In Proceedings of the Conference on Information Visualisation, pages 35–42, 2010.

[30] W. Katon. Asthma, Suicide Risk, and Psychiatric Comorbidity. American Journal of Psychiatry,167(9):1020–1022, 2010.

[31] R. A. Kerber. Method for calculating risk associated with family history of a disease. Genetic Epidemiology, 12(3):291–301, 1995.

[32] A. Kerren, H. C. Purchase, and M. Ward, editors. Multivariate Network Visualization. Number 8380 in Lecture notes in computer science. Springer, 2014.

[33] E. Kerzner, A. Lex, C. L. Sigulinsky, R. E. Marc, B. W. Jones, T. Urness, and M. Meyer. Graffinity: Visualizing Connectivity in Large Graphs. Computer Graphics Forum (EuroVis ’17), 36(3):251–260, 2017.

[34] N. W. Kim, S. K. Card, and J. Heer. Tracing Genealogical Data with TimeNets. In Proceedings of the International Conference on Advanced Visual Interfaces, AVI ’10, pages 241–248. ACM, 2010.

[35] S. Knight, R. P. Abo, H. J. Abel, D. W. Neklason, T. M. Tuohy, R. W. Burt, A. Thomas, and N. J. Camp. Shared Genomic Segment Analysis: The Power to Find Rare Disease Variants. Annals of Human Genetics,76(6):500–509, 2012.

[36] J. B. Kruskal and J. M. Landwehr. Icicle Plots: Better Displays for Hierarchical Clustering. The American Statistician,37(2):162, 1983.

[37] H. Lam, E. Bertini, P. Isenberg, C. Plaisant, and S. Carpendale. Empirical Studies in Information Visualization: Seven Scenarios. IEEE Transactions on Visualization and Computer Graphics,18(9):1520–1536, 2012.

[38] B. Lee, L. Nachmanson, G. Robertson, J. M. Carlson, and D. Heckerman. PhyloDet: A scalable visualization tool for mapping multiple traits to large evolutionary trees. Bioinformatics,25(19):2611–2612, 2009.

[39] B. Lee, L. Nachmanson, G. G. Robertson, J. Carlson, and D. Heckerman. Det.(distance encoded tree): A scalable visualization tool for mapping multiple traits to large evolutionary trees. MSR Tech Report MSR-TR- 2008-97, 2008.

[40] B. Lee, C. Parr, C. Plaisant, B. Bederson, V. Veksler, W. Gray, and C. Kotfila. TreePlus: Interactive Exploration of Networks with Enhanced Tree Layouts. IEEE Transactions on Visualization and Computer Graphics,12(6):1414–1426, 2006.

[41] B. Lee, C. Plaisant, C. S. Parr, J.-D. Fekete, and N. Henry. Task Taxonomy for Graph Visualization. In Proceedings of the AVI Workshop on BEyond Time and Errors: Novel Evaluation Methods for Information Visualization (BELIV ’06), pages 1–5, 2006.

[42] A. Lex, M. Streit, E. Kruijff, and D. Schmalstieg. Caleydo: Design and Evaluation of a Visual Analysis Framework for Gene Expression Data in its Biological Context. In Proceedings of the IEEE Symposium on Pacific Visualization (PacificVis ’10), pages 57–64. IEEE, 2010.

[43] A. Lex, M. Streit, C. Partl, K. Kashofer, and D. Schmalstieg. Comparative Analysis of Multidimensional, Quantitative Data. IEEE Transactions on Visualization and Computer Graphics (InfoVis ’10), 16(6):1027–1035, 2010.

[44] A. Lex, M. Streit,H.-J. Schulz, C. Partl, D. Schmalstieg, P. J. Park, and N. Gehlenborg. StratomeX: Visual Analysis of Large-Scale Heterogeneous Genomics Data for Cancer Subtype Characterization. Computer Graphics Forum (EuroVis ’12), 31(3):1175–1184, 2012.

[45] Y. Liu, S. Dai, C. Wang, Z. Zhou, and H. Qu. GenealogyVis: A System for Visual Analysis of Multidimensional Genealogical Data. IEEE Transactions on Human-Machine Systems, PP(99):1–13, 2017.

[46] V.-P. Mäkinen, M. Parkkonen, M. Wessman,P.-H. Groop, T. Kanninen, and K. Kaski. High-throughput pedigree drawing. European Journal of Human Genetics,13(8):987–989, May 2005.

[47] T. A. Manolio, F. S. Collins, N. J. Cox, D. B. Goldstein,L. A. Hindorff,D. J. Hunter, M. I. McCarthy, E. M. Ramos, L. R. Cardon, A. Chakravarti, J. H. Cho, A. E. Guttmacher, A. Kong, L. Kruglyak, E. Mardis, C. N. Rotimi, M. Slatkin, D. Valle, A. S. Whittemore, M. Boehnke, A. G. Clark,E. E. Eichler, G. Gibson, J. L. Haines, T. F. C. Mackay,S. A. McCarroll, and P. M. Visscher. Finding the missing heritability of complex diseases. Nature,461(7265):747–753, 2009.

[48] C. R. Marshall and the CNV and Schizophrenia Working Groups of the Psychiatric Genomics Consortium. Contribution of copy number variants to schizophrenia from a genome-wide study of 41,321 subjects. Nature Genetics, 49(1):27, 2017.

[49] A. Mazeika, J. Petersons, and M. H. Böhlen. PPPA: Push and Pull Pedigree Analyzer for Large and Complex Pedigree Databases. In Advances in Databases and Information Systems, pages 339–352, 2006.

[50] M. J. McGuffin and R. Balakrishnan. Interactive visualization of genealogical graphs. In Proceedings of the IEEE Symposium on Information Visualization (InfoVis ’05), pages 16–23, 2005.

[51] P. McGuffin, A. Marušič, and A. Farmer. What can psychiatric genetics offer suicidology? Crisis: The Journal of Crisis Intervention and Suicide Prevention, 22(2):61–65, 2001.

[52] M. Meyer, B. Wong, M. Styczynski, T. Munzner, and H. Pfister. Pathline: A Tool For Comparative Functional Genomics. Computer Graphics Forum (EuroVis ’10), 29(3):1043–1052, 2010.

[53] E. S. Mills. Genealogy in the information age: History’s new frontier. National Genealogical Society Quarterly, 91(91):260–277, 2003.

[54] E. T. Monson,K. de Klerk, S. C. Gaynor, A. H. Wagner, M. E. Breen, M. Parsons, T. L. Casavant, P. P. Zandi, J. B. Potash, and V. L. Willour. Whole-gene sequencing investigation of SAT1 in attempted suicide. American Journal of Medical Genetics Part B: Neuropsychiatric Genetics,171(6):888–895, 2016.

[55] T. Munzner, F. Guimbretière, S. Tasiran, L. Zhang, and Y. Zhou. TreeJuxtaposer: Scalable Tree Comparison Using Focus+Context with Guaranteed Visibility. In Proceedings of the ACM Conference on Computer Graphics and Interactive Techniques (SIGGRAPH ’03), pages 453–462. ACM, 2003.

[56] National Center for Health Statistics (US). Health, United States, 2015: With Special Feature on Racial and Ethnic Health Disparities. Health, United States. National Center for Health Statistics (US), 2016.

[57] R. A. Neher and T. Bedford. Nextflu: Real-time tracking of seasonal influenza virus evolution in humans. Bioinformatics,31(21):3546–3548, 2015.

[58] C. Partl, S. Gratzl, M. Streit, A. M. Wassermann, H. Pfister, D. Schmalstieg, and A. Lex. Pathfinder: Visual Analysis of Paths in Graphs. Computer Graphics Forum (EuroVis ’16), 35(3):71–80, 2016.

[59] C. Partl, A. Lex, M. Streit, D. Kalkofen, K. Kashofer, and D. Schmalstieg. enRoute: Dynamic Path Extraction from Biological Pathway Maps for Exploring Heterogeneous Experimental Datasets. BMC Bioinformatics, 14(Suppl 19):S3, 2013.

[60] T. Paterson, M. Graham, J. Kennedy, and A. Law. VIPER: A visualisation tool for exploring inheritance inconsistencies in genotyped pedigrees. BMC Bioinformatics,13(8):S5, 2012.

[61] N. L. Pedersen and A. Fiske. Genetic influences on suicide and nonfatal suicidal behavior: Twin study findings. European Psychiatry,25(5):264–267, 2010.

[62] Progeny Genetics LLC. Progeny, 2016.

[63] H.-J. Schulz. Treevis.net: A Tree Visualization Reference. IEEE Computer Graphics and Applications, 31(6):11–15, 2011.

[64] M. Sedlmair, M. Meyer, and T. Munzner. Design Study Methodology: Reflections from the Trenches and the Stacks. IEEE Transactions on Visualization and Computer Graphics,18(12):2431–2440, 2012.

[65] J. Seo and B. Shneiderman. Interactively Exploring Hierarchical Clustering Results. Computer,35(7):80–86, 2002.

[66] R. Shannon, T. Holland, and A. Quigley. Multivariate Graph Drawing using Parallel Coordinate Visualisations. Technical report, University of St Andrews, 2008.

[67] P. D. Shaw, M. Graham, J. Kennedy, I. Milne, and D. F. Marshall. Helium: Visualization of large scale plant pedigrees. BMC Bioinformatics, 15:259, 2014.

[68] M. Sokolowski, J. Wasserman, and D. Wasserman. Polygenic associations of neurodevelopmental genes in suicide attempt. Molecular Psychiatry, 2015.

[69] J. Stasko and E. Zhang. Focus+Context Display and Navigation Techniques for Enhancing Radial, Space-Filling Hierarchy Visualizations. In Proceedings of the IEEE Symposium on Information Vizualization (InfoVis ’00), pages 57–65. IEEE Computer Society Press, 2000.

[70] H. Thiele and P. Nürnberg. HaploPainter: A tool for drawing pedigrees with complex haplotypes. Bioinformatics, 21(8): 1730–1732, 2005.

[71] C. Tuttle, L. G. Nonato, and C. Silva. PedVis: A Structured, Space-Efficient Technique for Pedigree Visualization. IEEE Transactions on Visualization and Computer Graphics,16(6):1063–1072, 2010.

[72] S. van den Elzen and J. van Wijk. Multivariate Network Exploration and Presentation: From Detail to Overview via Selections and Aggregations. IEEE Transactions on Visualization and Computer Graphics (InfoVis ’14), 20(12):2310–2319, 2014.

[73] R. E. Voorrips, M. C. A. M. Bink, V. D. Weg, and W. Eric. Pedimap: Software for the Visualization of Genetic and Phenotypic Data in Pedigrees. Journal of Heredity,103(6):903–907, 2012.

[74] R. G. Waller, T. M. Darlington, X. Wei, M. Madsen, A. Thomas, K. Curtin, H. Coon, V. Rajamanickam, J. Musinsky, D. Jayabalan, D. Atanackovic, V. Rajkumar, S. Kumar, S. Slager, M. Middha, P. Galia, D. Demangel, M. Salama, V. Joseph, J. McKay, K. Offit, R. J. Klein, S. M. Lipkin, C. Dumontet, C. M. Vachon, and N. J. Camp. Novel pedigree analysis implicates DNA repair and chromatin remodeling in Multiple Myeloma risk. bioRxiv, page 137000, 2017.

[75] C. Ware, H. Purchase, L. Colpoys, and M. McGill. Cognitive Measurements of Graph Aesthetics. Information Visualization, 1(2):103–110, 2002.

[76] J. Xu, K. Kochanek, S. Murphy, and B. Tejada-Vera. National Vital Statistics Reports. Deaths: Final Data for 2007. Center for Disease Control and Prevention Division of Vital Statistics, 58, 2010.

